# Population genomics reveals multi-scale mechanisms sustaining schistosomiasis re-emergence in a near-elimination setting

**DOI:** 10.64898/2026.03.30.715205

**Authors:** Hannah Guss, Yannick Francioli, Elise N. Grover, Andrew Hill, Wei Zou, Kristen J. Wade, Hamish Pike, Siddharth S. Gopalan, Yang Liu, Bo Zhong, David D. Pollock, Elizabeth J. Carlton, Todd A. Castoe

**Author notes:** Corresponding author: Todd A. Castoe Department of Biology, University of Texas at Arlington, 501 S. Nedderman Drive, 337 Life Science; Arlington, TX 76010-0498.

## Abstract

In China, sustained snail control, environmental management, and mass drug administration with praziquantel reduced schistosomiasis to near-elimination levels, yet re-emergence in Sichuan Province during the early 2000s exposed vulnerabilities in late-stage control. We conducted a novel investigation of *Schistosoma japonicum* re-emergence in Sichuan in order to identify the processes underlying re-emergence and to showcase how genomic data can be used in such investigations. We sequenced whole genomes from 270 miracidia collected from 53 human hosts across 17 villages in 2007 – the year after re-emergence was documented.

Population genomic analyses revealed a broadly cohesive regional parasite population with weak geographic structuring. Genome-wide diversity was substantial, and demographic reconstructions revealed no recent decline in effective population size, demonstrating that parasite populations were not demographically fragmented at the onset of re-emergence, hinting at maintenance in reservoir populations. At finer spatial scales, several villages exhibited low diversity and elevated inbreeding, consistent with localized transmission maintained by small founding populations. Estimates of pairwise genetic relatedness revealed dense within-village sibling clusters alongside second- and third-degree relationships spanning villages, and rare first- and second-degree cross-village links. These findings are consistent with highly focal transmission with episodic parasite dispersal across villages, leading to a regional transmission network. Inferred minimum worm burden across hosts varied from 1 to 11 adult worm pairs, indicating heterogeneity in within-host parasite diversity, although uneven sampling limited inference. Together, these findings indicate that schistosomiasis re-emergence in this near-elimination setting was likely facilitated by a diverse parasite population maintained in reservoir populations, and that transmission, while predominantly local was occurring across a network of connected villages. This work illustrates how population genomics can reveal mechanisms driving re-emergence in late-stage elimination in complex, multi-host transmission systems.

**Author Summary:** Schistosomiasis is a parasitic disease that affects millions of people worldwide. Understanding the factors that contribute to persistence and transmission, even in the face of control programs, is central to reducing and ultimately eliminating the disease. In China, decades of intensive control efforts have dramatically reduced infections, yet the disease continued to persist in some regions. In this study, we analyzed the genomes of 270 parasite larvae collected from infected humans across 17 villages in Sichuan Province, China, shortly after schistosomiasis re-emerged in the region. By examining genetic relationships among parasites, we reconstructed patterns of transmission across villages and within individual infections. Our findings suggest that, despite major reductions in human infection prevalence, parasite populations sizes do not show signatures of recent population decline, indicating that non-human hosts may serve as key reservoirs promoting persistence. We also find that local transmission pathways, including clonal infections infecting multiple humans derived from a relatively small number of snail hosts, maintain parasite transmission pathways. Collectively, these findings suggest non-human hosts, intact local transmission pathways, and regional connectivity together enabled re-emergence, highlighting important challenges for elimination programs

## Introduction

Schistosomiasis is a neglected tropical disease that has been targeted for elimination as a public health problem by the World Health Organization[1], but the re-emergence and persistence of infections in transmission hotspots are a key barrier to attaining that goal [2,3]. Schistosomiasis is caused by parasitic blood flukes of the genus *Schistosoma* that affects more than 250 million people worldwide and contributes substantially to chronic morbidity, including hepatic fibrosis and anemia [4,5]. Among human schistosomes, *S. japonicum* is distinguished by its broad mammalian host range and zoonotic transmission ecology, infecting humans as well as bovines and other domestic and wild mammals [6–8]. Over the past seven decades, China has implemented one of the most sustained and comprehensive schistosomiasis control programs globally, combining large-scale snail control, environmental modification, health education, and mass drug administration (MDA) with praziquantel (PZQ) [9–11]. These coordinated interventions substantially reduced prevalence in many regions and brought transmission to near-elimination levels [2,12]. From both epidemiological and theoretical perspectives, such sustained reductions in prevalence are expected to push parasite populations toward transmission breakpoints and eventual collapse.

Classical macroparasite transmission theory predicts that sustained reductions in worm burden decrease mating probability and can ultimately drive parasite populations below density-dependent transmission breakpoints [13–15]. Because schistosomes are sexually reproducing and require male-female pairing within hosts, reductions in worm density can generate nonlinear declines in reproductive success. Under such models, declining human prevalence that goes below approximately 1% is expected to approach thresholds at which transmission becomes unsustainable in the absence of continued reintroduction via spatial connectivity or alternative host reservoirs [14]. Yet in the early 2000s, schistosomiasis re-emerged in mountainous regions of Sichuan Province despite prevalence having previously been estimated to be below this threshold [2]. This re-emergence challenges assumptions about how transmission collapse unfolds in advanced control settings.

Several non-mutually exclusive mechanisms could explain the persistence and/or re-emergence of transmission near elimination. First, effective population size (N_e_) may remain sufficiently large to buffer demographic collapse despite low observed human prevalence, particularly if parasite populations are harbored across multiple reservoir host species. In multi-host systems, reductions in human prevalence may not translate directly into proportional reductions in parasite population size if non-human reservoirs maintain transmission cycles [16–18]. Second, focal transmission may persist locally due to the survival of intermediate snail hosts or pockets of viable snail habitat that permit small founder populations to expand [7,10]. Third, regional connectivity via hydrological networks or host movement may prevent epidemiological fragmentation, allowing parasite lineages to reseed local populations even after substantial reductions in prevalence [19–21].

Disentangling these mechanisms require approaches that integrate demographic, spatial, and within-host scales of inference. Population genomics is such an approach that enables reconstruction of effective population size trajectories, quantification of genetic connectivity, and identification of close kin relationships among parasites [18,22–24]. Furthermore, noting that the cercarial life stage can be difficult to find and sample effectively, assessing schistosome populations through whole-genome sequencing of individual miracidia from fecal material permits inference of within-host parasite lineage structure without invasive sampling, providing a window into worm-pair number, diversity, and transmission heterogeneity. Population genomic inference can therefore reveal otherwise hidden mechanisms of persistence and connectivity not apparent from prevalence data alone.

Here, we leverage whole-genome sequencing of 270 *S. japonicum* miracidia collected from 53 human hosts across 17 villages in Sichuan Province in 2007, the year following documented re-emergence. We explicitly test key assumptions of density-dependent breakpoint models by evaluating whether sustained reductions in human prevalence were accompanied by detectable genomic evidence of parasite population decline, whether parasite populations became spatially fragmented consistent with disruption of local transmission networks, whether regional connectivity persisted at levels sufficient to reseed local populations, and whether within-host parasite structure revealed heterogeneity in worm burden that could enable a subset of hosts to disproportionately sustain transmission. By integrating genomic demographic inference with fine-scale relatedness analyses, we assess whether re-emergence reflects demographic collapse and episodic reseeding or instead sustained multi-scale transmission processes in which reductions in human prevalence did not translate into collapse of parasite populations.

## Materials and Methods

### Sample collection

270 *Schistosoma japonicum* miracidia were collected in 2007 from 53 human hosts across 17 villages in three counties of Sichuan Province, China, during village-wide infection surveys conducted as previously described [25]. Participants were screened for infection using the miracidia hatching test using stool samples collected over up to three consecutive days.

Miracidia emerging from positive hatching assays were individually isolated using a hematocrit tube or flame-drawn Pasteur pipette, washed three times in autoclaved deionized water, and transferred onto Whatman FTA® indicating cards (GE Healthcare) for long-term storage [26]. All sampling procedures were approved by the Sichuan Institutional Review Board, the University of California, Berkeley Committee for the Protection of Human Subjects, and the Colorado Multiple Institutional Review Board (protocol 15-1059). Written informed consent was obtained from all participants. Individuals who tested positive for *S. japonicum* infection were notified and referred to the local anti-schistosomiasis control station for treatment according to standard protocols. From the archived collection, we selected multiple miracidia per infected host across villages spanning approximately 75 km to maximize representation of regional genetic diversity. Not all infected hosts or villages are represented in the final dataset due to whole-genome amplification or sequencing failures, and because a subset of samples had been previously used for reduced-representation sequencing [27].

### Whole genome amplification and sequencing

Miracidia were recovered from Whatman FTA® cards using a Whatman Harris 2 mm micro-core punch (Whatman; cat. WB100029). Whole genome amplification (WGA) was performed directly from card punches following previously described protocols [27,28] using the Illustra Ready-To-Go GenomiPhi V3 DNA Amplification Kit (GE Healthcare; cat. 25-6601-96). Amplified DNA quantity was assessed using a Qubit fluorometer, and samples yielding >20 ng were selected for whole-genome shotgun library preparation. Sequencing libraries were prepared from individual amplified samples using Illumina Nextera Flex kits, pooled in multiplexed sets of up to 96 libraries, and sequenced on an Illumina NovaSeq 6000 S4 flow cell with 150 bp paired-end reads. Sequencing depth targeted approximately 20× genome coverage per miracidium. In total, 395 archival miracidia collected in 2007 and preserved on FTA cards underwent whole genome amplification, of which 270 yielded sufficient coverage and quality for downstream analyses.

### Read processing and variant identification

Across the 270 samples, the mean paired-end sequencing reads per sample was 75 million, the mean depth of coverage was 37x, and the mean percentage of reads that mapped to the reference *S. japonicum* genome [29] was 95% (S1. A and B). We quality filtered raw reads using Trimmomatic v.0.39 with the following options: LEADING:20 TRAILING:20 MINLEN:75 AVGQUAL:20 and then mapped our trimmed and quality filtered reads to the *S. japonicum* reference genome (ASM636876v1) using default parameters in BWA (Li and Durbin 2009). We called variants using GATK v.4.0.8.1 using the best practices workflow [30] by first generating individual variant call files (VCFs) using ‘HaplotypeCaller’, specifying –ERC GVCF and then called population variants using ‘GenotypeGVCFs’. We then used GATK ‘VariantFiltration’ to further hard filter based on GATK’s best practices recommendation (QD < 2, QUAL < 30.0, SOR > 3.0, FS > 60.0, MQ < 40.0, MQRankSum < -12.5, ReadPosRankSum < -8.0, cluster-size 3, cluster-window-size 10) and then used the module ‘SelectVariants’ to exclude non-filtered variants. We used VCFtools [31] to remove indels and retain biallelic sites that had a MAF > 0.05, genotype quality > 30 and removed sites that were missing more than 80% of genotype calls. After filtering variants for quality, completeness, and masking annotated genomic repeat elements, the final VCF contained 12,710,408 high quality single nucleotide polymorphisms (SNPs). For analyses sensitive to linkage disequilibrium or data density, we generated a reduced dataset by randomly selecting a single SNP per 10kb window, yielding 54,890 SNPs (hereafter referred to as the 10kb-thinned dataset). Because of the high proportion of repetitive DNA found in the *S*. *japonicum* genome (∼45%), and the difficulty in accurately calling variants in repeat regions, we also masked repetitive elements using the repeat annotation file from the *S. japonicum* reference genome (ASM636876v1) and removed all variants found on the Z chromosome.

### Population clustering and structure

We used the program ADMIXTURE v.1.3 [32] to estimate population structure using only our 10kb-thinned dataset. We used the cross-validation method implemented in ADMIXTURE to determine the best fit value of *K* from a range of values (*K=*2-10). To visually examine the distribution of genetic variance, we performed a principal components analysis (PCA) in PLINK v1.9 using our 10kb-thinned dataset including outgroup samples. We used R version 4.4.3 to calculate the percent of variance explained by each principal component and then plotted the first four principal components. Using the same dataset, we then inferred relationships among individuals by constructing a neighbor-joining tree from average pairwise distances using the R package ‘ape’ [33].

### Phylogenic analysis

To further visualize genetic relationships among individual miracidia, we constructed a phylogenetic tree using the same 10kb-thinned SNP dataset. Pairwise genetic distances among individuals were calculated from genome-wide SNP variation, producing a matrix of average genetic distances between all samples. Using this distance matrix, we inferred a neighbor-joining tree using the R package *ape* [33]. Sample metadata linking each miracidium to its village of origin were then incorporated for visualization, and the resulting phylogeny was plotted in R using the package *ggtree*. The tree was displayed in a circular (“fan”) layout with tips colored according to village identity to facilitate interpretation of spatial population structure.

### Estimating genomic diversity

To quantify genomic diversity across samples and villages, we estimated per-sample heterozygosity, genome-wide inbreeding coefficients (F), and nucleotide diversity (π) from the quality-filtered SNP dataset. Heterozygosity and inbreeding coefficients were calculated from individual-level genotype counts using the PLINK --het framework, with heterozygosity estimated as 1 ― (OHOM/𝑁_SITES). We summarized heterozygosity and F values for each miracidium and compared their distributions across villages. To estimate nucleotide diversity, we calculated genome-wide π in sliding windows and generated per-sample π estimates for comparison with individual inbreeding coefficients. We then used R version 4.4.3 to visualize variation in heterozygosity, F, and π across villages and geographic groups, and to test the relationships between F and both heterozygosity and per-sample π using linear models.

### Demographic inferences

To infer the demographic history of the Sichuan *S. japonicum* population, we first used easySFS [34] to generate a folded site frequency spectrum (SFS), with projection set to 540 to retain all sites across individuals, resulting in 6,157,643 segregating sites. We then ran Stairway Plot v2.1.1 [34] using the assembled genome size (344 Mb) as the sequence length, 540 chromosomes, and assuming a generation time of 0.5 years and a mutation rate of 8.9 × 10⁻⁹ per site per generation, consistent with published estimates for schistosomes [24]. We used default Stairway Plot parameters, including 200 bootstrap replicates and training on 67% of sites per replicate to obtain confidence intervals around the inferred demographic history. To further validate demographic inferences, we also applied SMC++ [35] using the same generation time and mutation rate parameters described above. We estimated population history over the interval of 10 to 100,000 generations, including all 270 samples and designating three as distinguished individuals. Variant data were split by chromosome with VCFtools [31] and converted to SMC input format for each distinguished individual using the vcf2smc command, with runs of homozygosity longer than 50 kb masked as missing. Demographic inference was then performed with the estimate command across all input files. To evaluate uncertainty, we conducted bootstrap analyses by resampling 10 randomly selected 10 Mb segments per chromosome.

### Inferences of fine-scale relatedness

To assess patterns of genetic relatedness across the parasite population, we used the RAB (Ratio of Allele Balance) metric implemented in NGSrelate v2 [36]. This method leverages genotype likelihoods to estimate pairwise relatedness and is particularly well suited for populations with high inbreeding. In this framework, RAB represents the probability that two individuals share alleles identical by descent (IBD) at a given locus accounting for inbreeding (Hedrick and Lacy 2015). We also used the Rare Allele sharing (RAS) pipeline [38,39], to infer RAS values between pairs of miracids. For both of these inferences, we first pruned the dataset using MAF >= 0.1 and missingness <= 0.2 filters, and pruned SNPs in strong linkage disequilibrium using PLINK v2.0 [40] by scanning the genome in sliding windows of 50 SNPs advancing by 10 SNPs and removing variants with pairwise r^2^ > 0.2 within each window. We visualized pairwise RAB values as heatmaps in R using ggplot2, with samples ordered by village in a predefined north-to-south sequence and, within villages, by host identity. Heatmap cells were colored according to either binned or continuous RAB values, and village boundaries were annotated along both axes to facilitate interpretation of within-host, within-village, and between-village patterns of relatedness.

Both RAB and RAS were further benchmarked against pedigree estimates reconstructed with the R package Sequoia v3.0.3 [41], which infers parent–offspring and sibship relationships via a maximum-likelihood framework and COLONY [42], another pedigree inference software.

Analyses in Sequoia and COLONY were conducted using the original the LD-pruned dataset used for RAS and RAB inferences, which was then further 100-kb–thinned (2,962 SNPs). This filtering approach followed recommendations in the documentation of these programs, and is required to accommodate the computational burden of pedigree reconstruction. We identified candidate related miracid pairs using the Sequoia command “GetMaybeRel”, the “ped” module with CalcLLR=TRUE, a conservative genotype error rate (Err=0.01), discrete generations, and summarized log-likelihood ratios (LLR) and assignment confidence (EstConf). For an independent cross-check, we inferred sibships with COLONY, treating loci as diploid/dioecious, allowing polygamy for both sexes and inbreeding, and assigning conservative genotyping error rates (allelic drop-out and other error = 0.01).

To convert continuous RAB values into categorical relatedness classes, we constructed empirical training sets for 1^st^ degree and unrelated (defined as 4th+ degree) relationships and then fit a simple normal-mixture framework to generate posteriors of relatedness degrees for every miracid pair. First, we defined a high-confidence full-sib set by intersecting Sequoia full-sib calls with COLONY full-sib dyads. We then defined an “unrelated” set by selecting pairs classified as unrelated by both methods and sampled from geographically distant villages (pairs between northern and southern groups). From these two reference sets, we estimated the empirical mean and variance of RAB for full siblings and unrelated pairs. Expected RAB means (𝜇_degree_) and variance (𝜎_degree_) for intermediate degrees of relatedness were then approximated by placing successive means halfway between the sibling and unrelated expectations, and by reducing the sibling variance by half for each additional degree of separation, reflecting the increased number of meiosis events 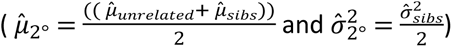. These parameters defined a four-component normal mixture corresponding to 1st-, 2nd-, 3rd-degree, and unrelated. Assuming equal prior probabilities across classes, we computed posterior probabilities for each category for every observed pairwise RAB value. For downstream analyses, we retained and summarized pairs whose maximum a posteriori (MAP) classification exceeded 0.95 posterior probability. Based on this posterior calibration, RAB thresholds corresponding to ≥95% posterior probability were defined as follows: first-degree (full sibling or parent–offspring) relationships were assigned at RAB > 0.425; second-degree relationships corresponded to RAB values between 0.231 and 0.425; third-degree relationships were defined as RAB values between 0.0166 and 0.231; and values < 0.0166 were classified as fourth-degree or unrelated. These calibrated thresholds were used consistently in downstream host-, village-, and network-level analyses, as well as in figure visualizations where RAB cutoffs are applied.

To visualize relatedness patterns, we generated network and circular (circos) representations from pairs classified with ≥0.95 posterior probability. Full-sibling clusters within hosts were defined using the calibrated first-degree RAB threshold (RAB > 0.425), and for cluster-level visualizations, multiple dyadic links between the same pair of clusters were collapsed to a single connection, retaining the highest relatedness class observed. Network plots were constructed in R v4.4.3 using the packages *igraph* and *ggraph*, applying a Fruchterman–Reingold force-directed layout. Circos visualizations were generated using the *circlize* package, with villages ordered north–south; host-level tiles represent individual hosts and cluster-level tiles represent inferred sibling clusters, with stacked points (and tile height in cluster panels) reflecting the number of sequenced miracidia. Links were drawn between hosts or clusters, colored by relatedness degree and scaled by the number of contributing dyads, and an outer track denotes geographic village groupings.

## Results

### Weak regional genetic structure supports cohesive demographic inference

The 270 whole genome amplified miracidia included in this analysis came from 53 human hosts and 17 villages dispersed across four geographic regions: North (N=5), Central (N=5), South Central (N=4) and South (N=3) (Fig 1A and S1 Table). Our PCA on the full SNP dataset (12,710,408 SNPs) indicated that samples from most villages clustered into a single large population (based on PC1-2), with limited differentiation except for villages on the northern and southern periphery (Fig 1B). Samples from village V, a geographically distant northern village (Fig 1A), formed a distinct cluster, while a subset of samples from the southernmost villages B and I also separated along the same axes (Fig 1B). Analysis of our 10 kb–thinned SNP dataset using ADMIXTURE (excluding all closely related individuals ( 1^st^ and most 2^nd^ degree with RAB < 0.3; 54,890 SNPs) suggested K = 1 as the optimal number of genetic clusters, while K = 2 captured a weak north–south gradient (S2). Together, PCA and ADMIXTURE support the presence of a largely cohesive regional population with modest north–south differentiation.

**Fig 1.**
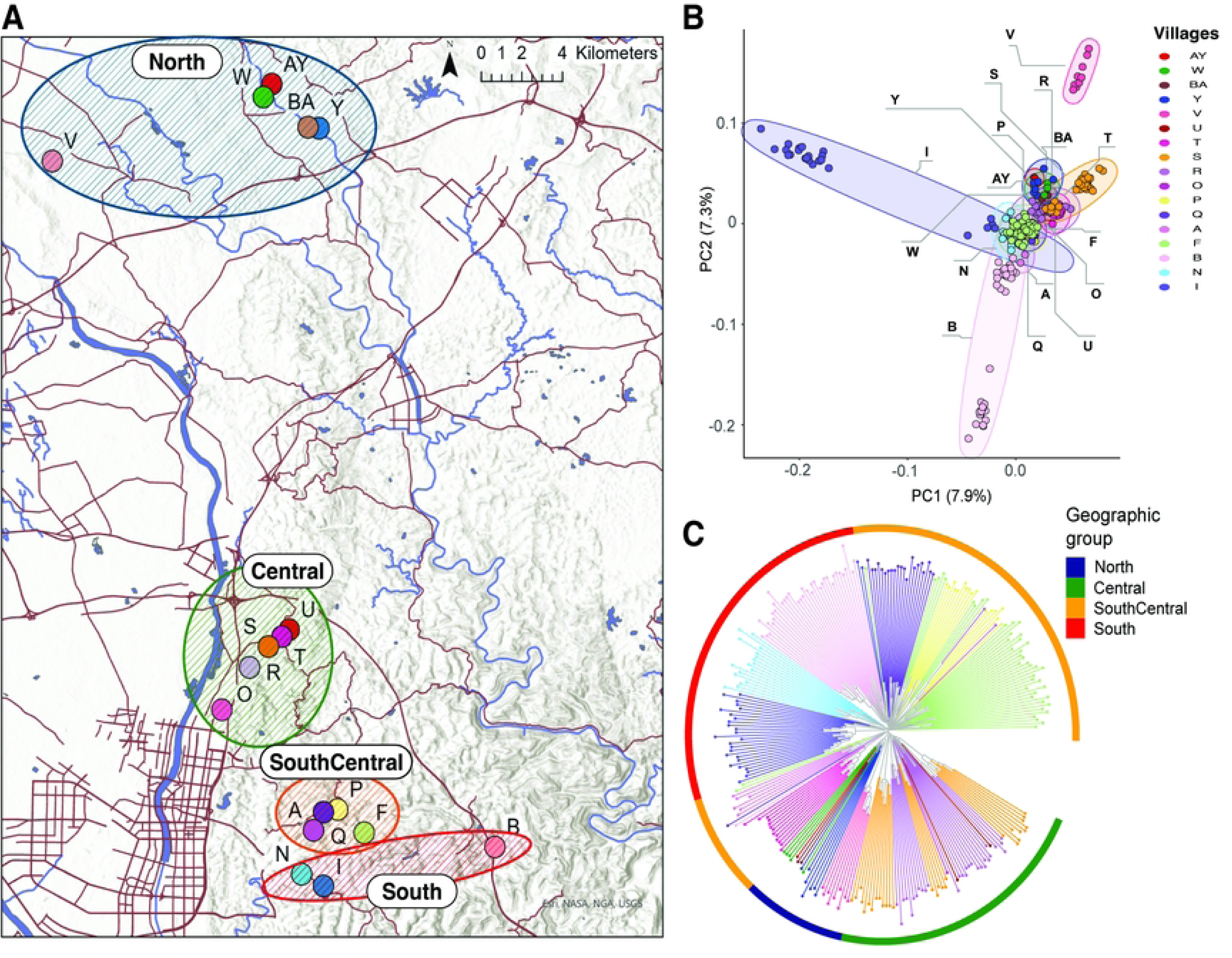
Geographic sampling and broad population genomic structure of Schistosoma japonicum miracidia in Sichuan Province, China. (A) Topographic map showing the locations of 17 sampled villages. Colored dots denote villages, with major waterways (blue) and roads (brown) providing geographic context. Geographic grouping is labeled with ovals. (B) Principal component analysis of genome-wide SNP variation among 270 miracidia. Most samples cluster within a single major population, whereas parasites from geographically peripheral northern (e.g., V) and southern (e.g., B and I) villages form more distinct groupings, indicating weak spatial substructure. (C) Neighbor-joining tree based on pairwise genome-wide genetic distances from a 10-kb thinned SNP dataset. Branch lengths represent relative genetic distance and were scaled for visualization. Branches are colored by village, revealing fine-scale clustering within villages and regional grouping of geographically proximate villages, alongside incomplete separation among several central villages consistent with shared ancestry and ongoing connectivity.

Phylogenetic analysis of the same 10kb-thinned dataset revealed finer-scale structuring within this broader regional population (Fig 1C). Samples from individual villages frequently clustered together, and geographically proximate villages often formed regional groupings. For example, the three southernmost villages (B, N, and I) grouped together, with samples clustering exclusively within their respective villages; a similar pattern of nearly exclusive within village clustering was observed among the northernmost villages (AY, W, BA, and Y; Fig 1C). In contrast, samples from central and south villages (e.g., S, F, P, and R) were less differentiated by village, with adjacent villages interspersed on the tree with one another, indicating incomplete geographic segregation and evidence of shared ancestry across these communities (Fig 1C).

Collectively, results from PCA, ADMIXTURE, and phylogenetic analyses indicate weak regional genetic structure superimposed on finer-scale local clustering of within villages and among adjacent villages.

### Genomic diversity

Given the intensive control of schistosome in this region – including multiple prior rounds of mass drug administration and improvements in sanitation – we expected parasite populations here to have reduced genomic diversity. In addition, the clonal amplification inherent to the schistosome life cycle could further elevate levels of inbreeding under conditions of restricted transmission. To address this, we estimated genome-wide inbreeding coefficients (F) for each miracidium and, contrary to expectation, observed wide variation across samples (F = –0.16 to 0.74) and among villages (mean F values 0.025–0.579), indicating substantial heterogeneity in inbreeding levels (Fig 2A). Villages A (𝐹 = 0.58), V (𝐹 = 0.58), and Y (𝐹= 0.54) had high F values, consistent with relatively restricted or locally isolated parasite populations and possible founder effects. In contrast, villages R (𝐹 = 0.39), P (𝐹 = 0.40), and N (𝐹 = 0.42) showed lower means and broader distributions (especially R and N) in F values, consistent with slightly greater genetic diversity and indicative of potentially more heterogeneous source populations (Fig 2A).

**Fig 2.**
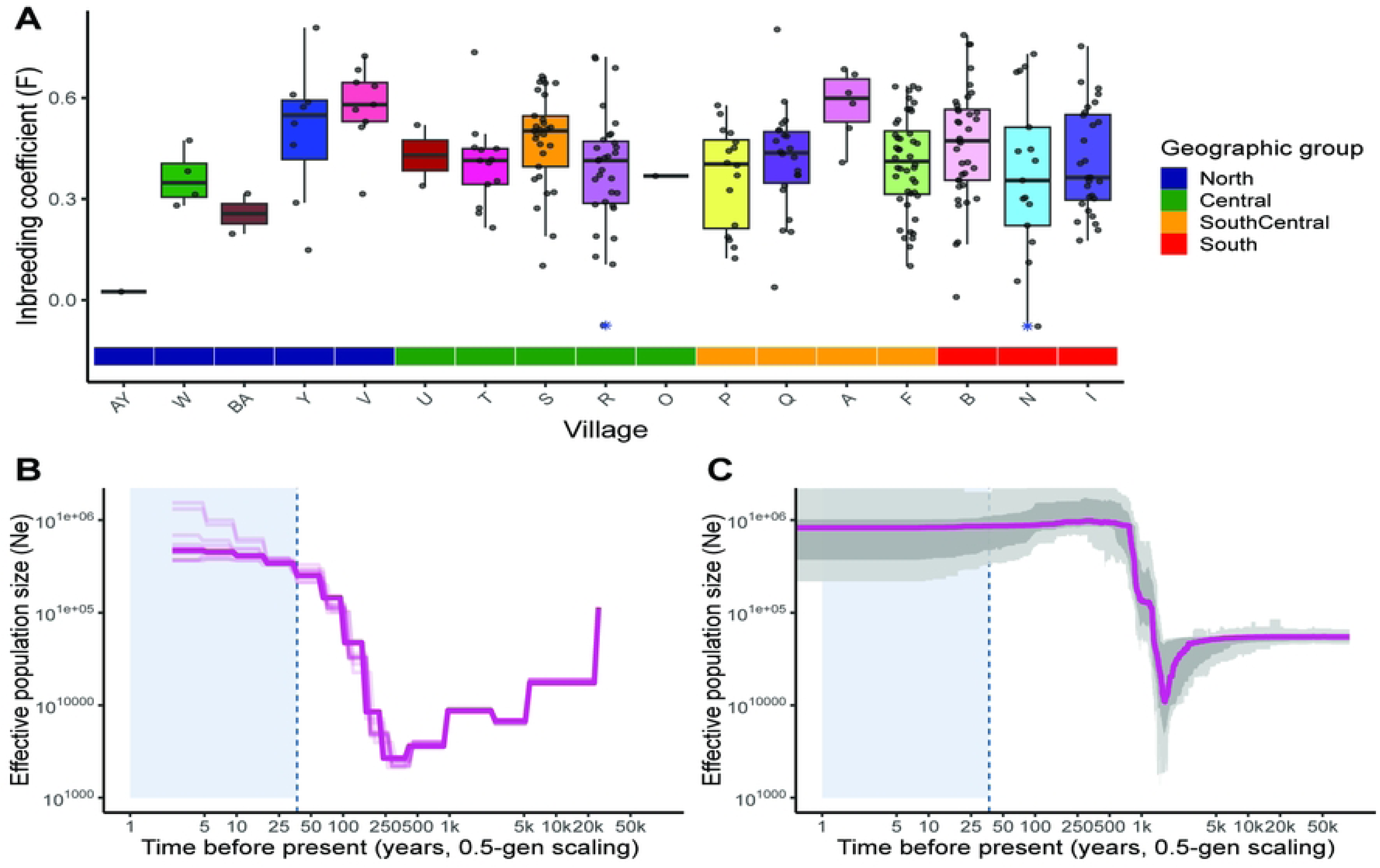
**Genomic diversity, inbreeding, and historical demography of Schistosoma japonicum in Sichuan Province**. (A) Genome-wide inbreeding coefficients (F) for individual miracidia summarized by village. Points represent individuals and boxplots show village-level distributions, colored by geographic region (B) Demographic history inferred using the sequential Markov coalescent approach (SMC++), showing a concordant deep historical decline in Ne without evidence of a strong recent reduction during the period of modern schistosomiasis control (demarcated by the blue dotted line and shaded region). (C) Historical effective population size (Ne) inferred from site-frequency spectra using Stairway Plot. The trajectory indicates a pronounced long-term population decline beginning approximately 1,500 years before present, followed by partial recovery and stabilization. Together, these results suggest that contemporary control efforts have not substantially reduced parasite effective population size, consistent with persistence maintained through non-human reservoir hosts.

Because the inbreeding coefficient (F) is calculated directly from individual heterozygosity relative to expected heterozygosity, a strong inverse relationship between F and heterozygosity (R² ≈ 1) is expected. By comparison, the observation of a negative correlation between F and nucleotide diversity (π) (R² = 0.72, p < 0.05) reflects a broader biologically informative association between individual-level inbreeding and genome-wide diversity across villages.

Together, these patterns reveal pronounced spatial heterogeneity in genomic diversity, with evidence of restricted transmission and elevated inbreeding in some villages yet maintained diversity in others. Notably, the persistence of moderate to high diversity in multiple villages is inconsistent with a uniform regional genetic bottleneck and instead indicates that parasite populations were not uniformly demographically collapsed prior to re-emergence.

### No genomic evidence of recent demographic collapse

To further evaluate whether the heterogeneous patterns of genomic diversity observed across villages reflect a recent regional demographic contraction, as opposed to village-specific founder effects or spatially restricted transmission, we leveraged genome-wide variation to infer historical changes in parasite effective population size (N_e_) using two complementary approaches: a site frequency spectrum–based method (Stairway Plot) and a sequential Markov coalescent framework (SMC++). Both analyses inferred a pronounced decline in N_e_ centered around approximately 1,500 years ago, followed by partial recovery and subsequent stabilization Figs. 2B-C). In both models, this historical decline substantially predates modern schistosomiasis control efforts initiated in the 1970s [10] and likely reflects older environmental or sociopolitical changes, such as shifts in land use, climate, or human settlement. Across both approaches, neither detected a recent, obvious decline in effective population size during the decades of intensive control that dramatically reduced human infection prevalence. Instead, inferred N_e_ remained relatively stable throughout the period corresponding to large-scale snail control and sustained mass drug administration. Thus, despite well-documented declines in human prevalence over this period, we find no genomic evidence of recent demographic collapse in parasite populations, suggesting that transmission was maintained through processes not fully captured by human case counts, including the likely contribution of non-human host reservoirs.

### Fine-scale relatedness reveals focal transmission within villages

To characterize fine-scale transmission dynamics within Sichuan, we compared RAB and RAS IBD approaches, each benchmarked against pedigree-based reconstructions generated using Sequoia and Colony. Although RAS and RAB were positively correlated (R² = 0.388), RAB more accurately recapitulated pedigree-based expectations: full-sibling pairs independently identified by Sequoia and Colony clustered near the expected IBD value of ∼0.5 under RAB, whereas RAS showed greater dispersion and inflation among unrelated pairs (S4. A–C). We therefore used RAB as the primary estimator of relatedness and calibrated posterior probability thresholds for relationship classes based on full-sibling pairs (from a single host) identified by both pedigree methods (mean RAB = 0.566; variance = 0.0031; n = 262 sibling pairs; see Methods).

Closely related parasites were common within individual hosts (n = 53): 70% of hosts harbored at least one first-degree (full-sibling) miracidial pair and 80% harbored at least one second-degree pair (Fig 3A and B). The frequent co-occurrence of half-siblings within hosts is consistent with repeated exposure to clonal cercariae originating from single infected snails, which produce genetically identical cercariae during asexual amplification. Considering the frequency of half-sib clusters, some full-sibling clusters may also represent clonal pairs. These within-host patterns are visually apparent in the RAB heatmap (Fig 4), where within-host sibling groups form distinct triangular blocks of high relatedness along the diagonal.

**Fig 3.**
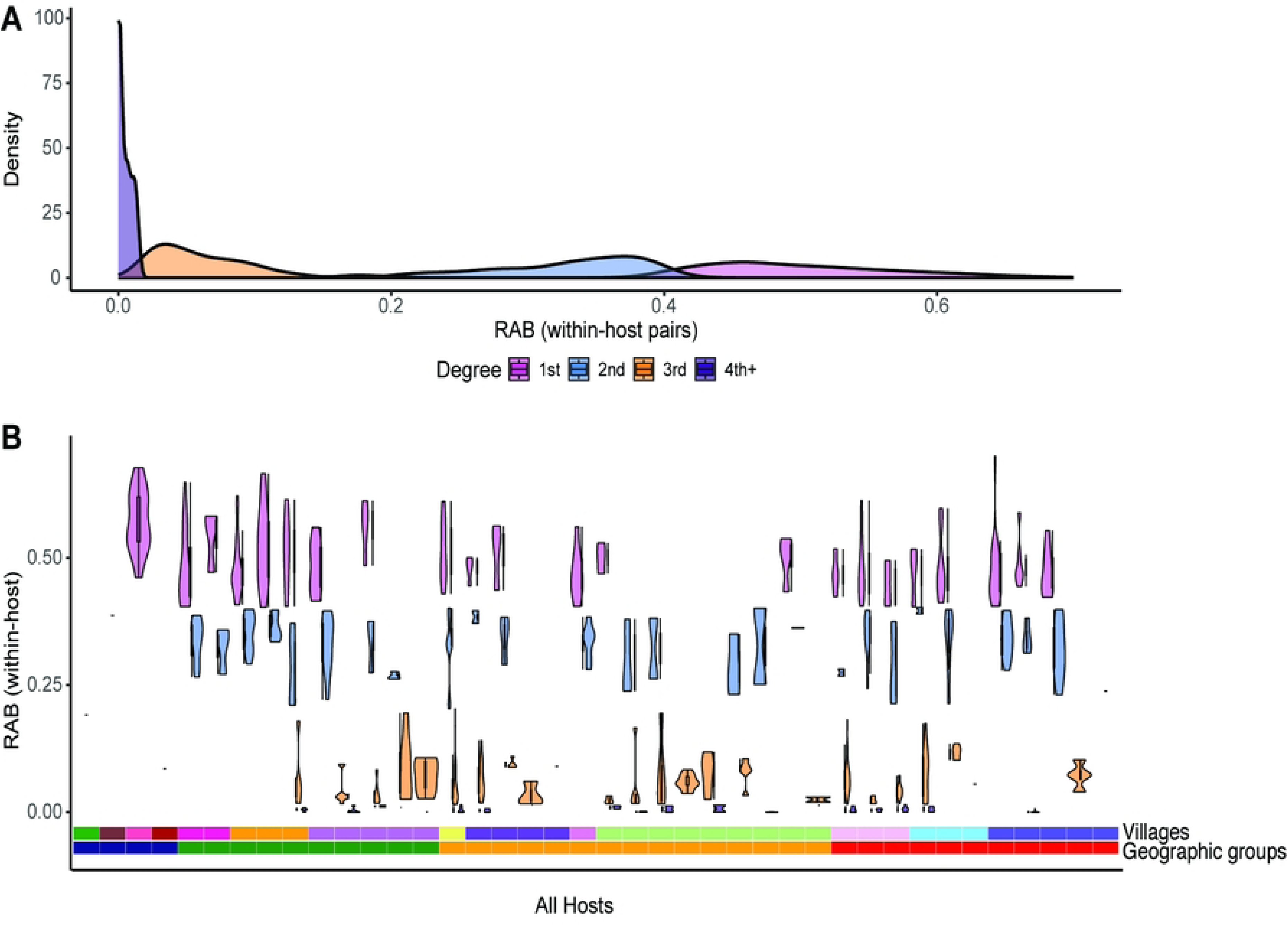
**Within-host distribution of RAB-inferred relatedness among Schistosoma japonicum miracidia**. (A) Density of Rare Allele Balance (RAB) values for miracidial pairs within hosts, colored by pedigree-calibrated relatedness class. (B) Host-specific distributions of within-host RAB values across all sampled hosts, with bars indicating village membership. Peaks of first- and second-degree relatedness reflect clonal infections derived from single snails, whereas substantial third-degree relatedness indicates localized inbreeding. These patterns demonstrate that clonality and inbreeding jointly structure infection within hosts and villages.

**Fig 4.**
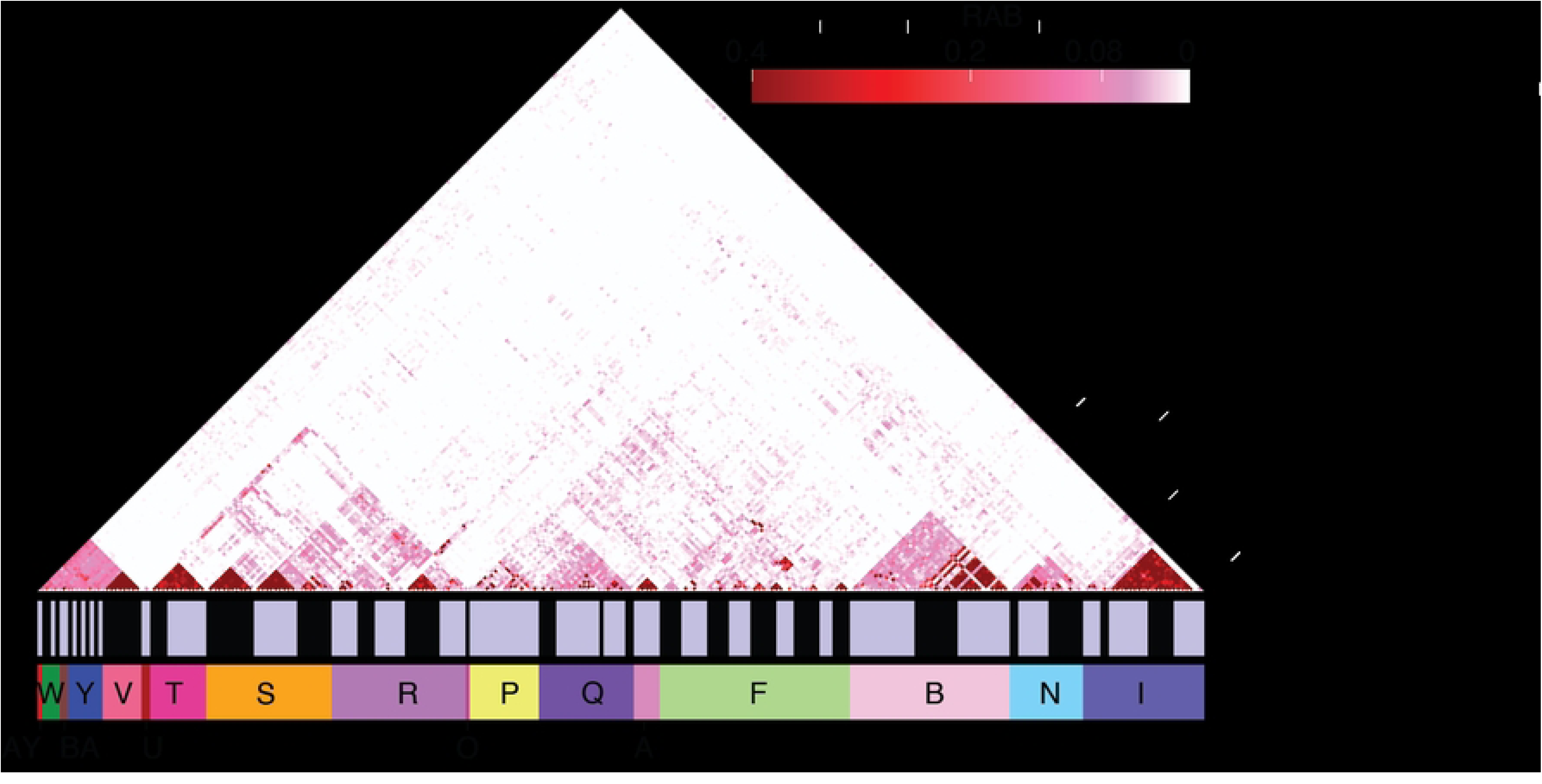
Genome-wide relatedness among Schistosoma japonicum miracidia. Triangle heatmap of pairwise allele-sharing (RAB) across all samples, ordered by village along a north–south gradient with hosts marked by alternating bars. High-RAB triangles (> 0.4) indicate within-host sibling groups, whereas broader regions of moderate relatedness (0.1–0.4) reflect relationships shared among hosts within villages and geographic clusters. Although relatedness shows hierarchical spatial structure, several first- and second-degree connections span distant villages (e.g., R, P, and U), indicating transmission links not explained solely by geographic distance.

At the village scale, the heatmap reveals larger triangular blocks of moderate relatedness among hosts within the same village (Fig 4), including first- and second-degree relationships shared across individuals. We observed clusters of first- and second-degree related miracidia infecting different individuals within villages, consistent with multiple hosts sharing exposure to the same infected snails or closely related parasite lineages circulating locally. In some villages (e.g., S and I), infections were composed almost exclusively of first- and second-degree related parasites, producing triangular blocks indicative of transmission maintained by a small number of parasite lineages and limited snail infection sources. In other villages, broader within-village relatedness distributions indicate contributions from a more heterogeneous local parasite and snail pool.

Together, the within host half-sibling relationships and within-village relatedness blocks demonstrates that village-level transmission frequently reflects localized amplification from limited numbers of infected snails. These dense local clusters of closely related parasites persist despite low overall regional prevalence, indicating that most transmission within villages is highly localized from limited, closely related infection sources.

### Cross-village relatedness indicates limited but persistent regional connectivity

Although closely related parasites were predominantly found within villages, we identified rare but notable instances of first- and second-degree relationships spanning villages. At the host level (Fig 5A), visualization of ≥95% posterior first- and second-degree links shows that nearly all (96.6%) close-kin connections occur among hosts within the same village. Eight first-degree clusters and three second-degree host pairs were detected, with only two connections linking different villages. One notable exception involved a host from village R that shared a first-degree relationship with a host in village P and a second-degree relationship with a host in village U. Apart from this cluster, close-kin relatedness appears highly localized, indicating that recent direct transmission between villages was uncommon. Visualization at the sibling-cluster level (Fig 5B) provides additional resolution. Rather than grouping links by host, this organizes connections by genetically inferred sibling clusters within hosts, which approximate distinct reproducing worm pairs within hosts. This representation similarly reveals that a subset of sibling clusters contributes disproportionately to between-host genetic links. Of the 152 identified sibling clusters, 33 had a first- or second-degree relationship with a sibling cluster from another host within the village, and 7 had a first- or second-degree relationship with a sibling cluster from another village. To further resolve the spatial topology of close-kin connections, we visualized first- and second-degree relationships as an individual-level network (Fig 6A). Consistent with the host- and sib-cluster–level patterns, close-kin relationships form tightly clustered, village-contained components, with only rare cross-village links. Notably, the few between-village connections are restricted to specific village pairs (R–P and R–U), reinforcing the conclusion that recent transmission is highly localized and only occasionally spans village boundaries.

**Fig 5.**
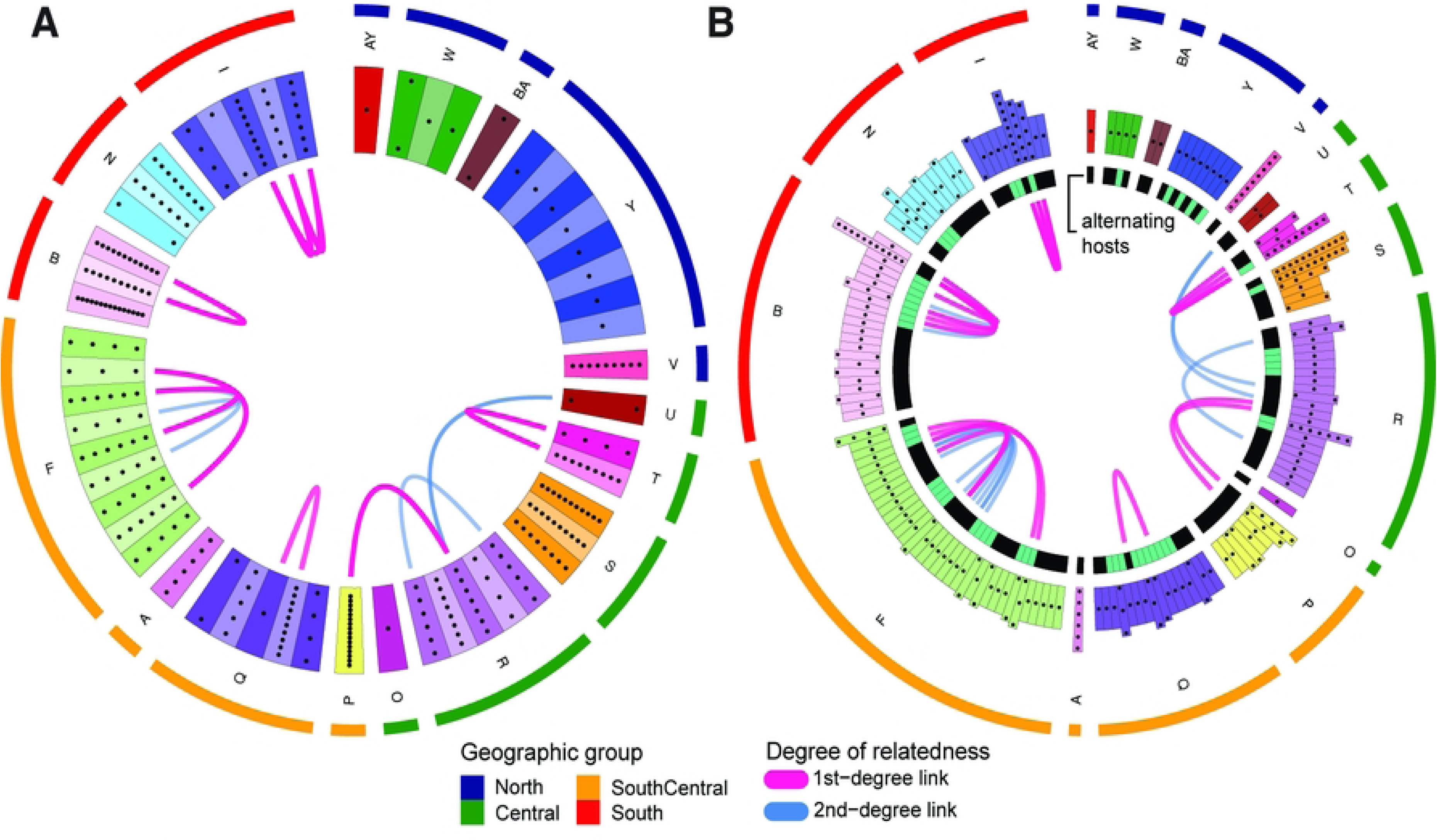
Host- and sib-cluster–level organization of close-kin relationships among *Schistosoma japonicum* miracidia in Sichuan Province, China. (A) Circos visualization of first- and second-degree relationships grouped by human host. Tiles represent hosts, dots denote sampled miracidia, and arcs indicate kinship (pink = first degree; blue = second degree). Close-kin links frequently connect multiple hosts within the same village, consistent with infection by clonal cercariae derived from few infected snails. Eight first-degree clusters and four second-degree host pairs were detected, with nearly all links connected within villages and only rare connections spanning villages. (B) Visualization at the sib-cluster level shows the distribution of genetically inferred sibling groups and variation in inferred worm burden. Inner bars represent sibling clusters with alternating host membership, and dots again denote individual miracidia. Hosts with higher worm burden more often participate in between-host kinship links, indicating that a limited number of reproducing worm pairs can sustain local transmission. Outer colored arcs denote geographic clusters.

**Fig 6.**
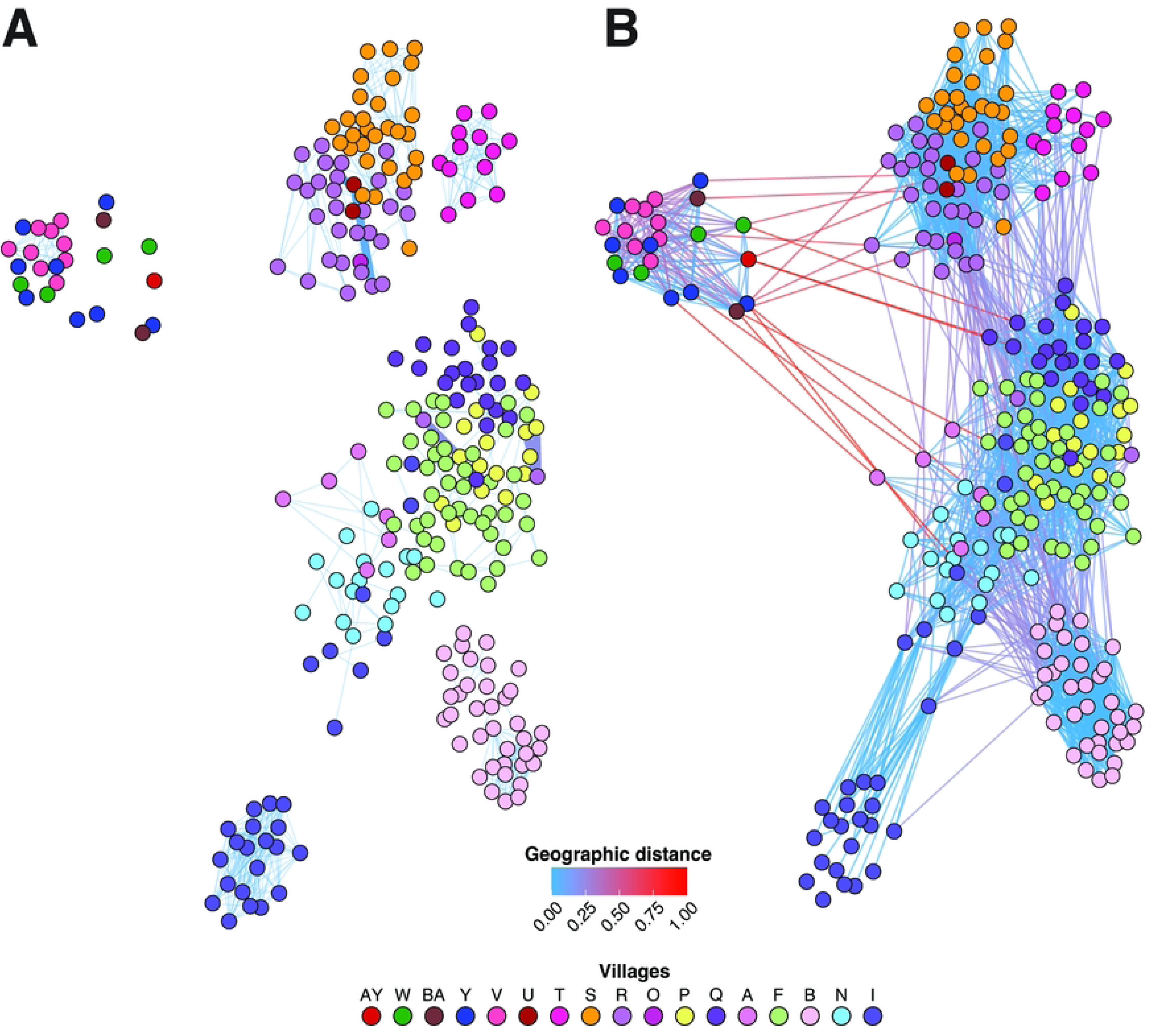
Network structure of close-kin and third-degree relationships among *S. japonicum* miracidia. Nodes represent individual miracidia and are colored by village of origin (north–south order shown in legend). Edges connect pairs of individuals inferred to be related with ≥95% posterior probability. Edge color reflects rescaled geographic distance between villages (blue = short distance; red = long distance). (A) First- and second-degree relationships (close kin) form largely village-contained clusters, with only rare cross-village links. Highlighted red connections indicate close-kin links between villages R–P and R–U. (B) Third-degree relationships reveal a substantially broader and more continuous spatial network, frequently connecting adjacent villages and extending across regional groupings (North, Central, Southcentral, South).

These network-based observations are consistent with patterns visible in the RAB heatmap introduced earlier (Fig 4). In addition to the prominent diagonal village-level blocks, the RAB heatmap reveals sparse but distinct off-diagonal hotspots of elevated relatedness connecting hosts from different villages. While first- and second-degree cross-village links are rare, moderate off-diagonal signals – corresponding primarily to second- and third-degree relationships – are more widespread, particularly within the central geographic cluster. This matrix-based view reinforces the conclusion that parasite relatedness is not structured strictly by geographic distance, as several geographically separated villages share detectable genetic connections.

In contrast to the rarity of close-kin cross-village links, third-degree relationships reveal a broader and more continuous spatial network (Fig 6B). Numerous third-degree links connect hosts across villages, frequently uniting geographically adjacent villages and, in some cases, spanning larger regional groupings. While certain villages remain relatively self-contained (e.g., village B), most are part of multi-village networks of relatedness structured largely by the four geographic village groups (North, Central, Southcentral, and South). Importantly, this broader network mirrors the weak north–south differentiation observed in PCA and ADMIXTURE analyses, reinforcing the conclusion that parasite populations across villages were not fully isolated. Together with the absence of recent demographic collapse, these results indicate that village-level transmission occurred recently (relative to our sampling) within a regionally connected parasite population.

### Heterogeneity in inferred worm burden and number of infection events

We used the sibling level clustering to infer the minimum number of unique infection events that contributed to infections within a given host, which provides an estimate of the intensity and heterogeneity of exposure in this population. We estimated the minimum number of reproducing worm pairs per host by counting distinct sibling clusters. Because sampling depth varied across hosts, worm-burden analyses were conducted with explicit consideration of sampling effort. We assumed each sibling cluster within a host represented a unique adult worm pair and, at a minimum, two successful infection events, one for each worm. We note that it is possible for a host to be infected with multiple clonal cercariae from a single snail (snails typically can tolerate infection by only one *S. japonicum* miracidia) however exposure to at least two infected snails is required to produce a reproducing worm pair and if such an exposure yielded multiple clonal worm pairs, it would be correctly counted as two unique infection events by our approach.

Among the 23 hosts with five or more miracidia sampled, inferred minimum worm burden ranged from 1 to 11 distinct sibling clusters, equivalent to 2 to 22 reproducing adult worms and unique infection events (S5), with substantial heterogeneity observed across villages. For example, villages B and R exhibited consistently higher inferred worm burdens (B: mean = 7, var = 16; R: mean = 4.6, var = 1.8), whereas village I showed consistently lower burdens (I: mean = 2.6, var = 0.8). Notably, among the seven hosts with >9 sampled miracidia, village S included hosts with 10–11 sampled miracidia that were all inferred to derive from a single parental worm pair, whereas hosts from villages P, B, and R harbored between 6–11 distinct sibling clusters (Supp. Fig 5). These contrasts indicate that substantial heterogeneity exists in worm burden across hosts and villages. These findings indicate some hosts have become infected via 20+ infection events, whereas others have far more limited exposure.

## Discussion

### Multi-scale genomic evidence of persistence in a near-elimination setting

In this retrospective genomic analysis of parasites collected immediately following documented re-emergence, we identified potential drivers of persistence across multiple biological and spatial scales. Parasites formed a broadly cohesive regional population with weak geographic structure. Despite decades of intensive control and marked reductions in human prevalence, genome-wide diversity remained high and demographic reconstructions revealed no evidence of recent effective population size decline. For example, standing genetic diversity in Sichuan miracidia (π ≈ 0.0015) was approximately half that reported for East African *S. mansoni* populations at much earlier stages of control (π ≈ 0.0033) [24]. Consistent with this inference of relatively high genetic diversity, our demographic reconstructions indicate that parasite effective population size has remained relatively large and stable despite prolonged control in Sichuan. At finer scales, within hosts and villages, dense clusters of closely related parasites within hosts and villages reflected clonal amplification from infected snails. Although first- and second-degree relationships were largely confined to within villages comparisons, third-degree connectivity indicated low-frequency but persistent genetic exchange among villages. Inferred worm burdens varied substantially among hosts, and hosts harboring more sibling clusters had more local relatives. Together, these findings indicate that re-emergence did not result from reintroduction into demographically collapsed parasite populations. Instead, transmission persisted through local amplification of limited numbers of genotypes embedded within a regionally connected and demographically stable parasite population.

Schistosomiasis elimination programs assume that sustained reductions in human prevalence will eventually drive parasite populations below density-dependent transmission breakpoints, leading to demographic collapse [13–15]. Both Stairway Plot and SMC++ detected no genomic evidence of a recent decline in effective population size corresponding to modern schistosomiasis control. Instead, both inferred an older contraction approximately 1,500 years ago followed by long-term stabilization. The absence of a recent bottleneck is notable given substantial reductions in human prevalence following sustained MDA, snail control, and environmental management [9,10]. Under these expectations, prolonged prevalence reductions should leave a genomic signature of declining effective population size. Our results from Sichuan reveal a more complex relationship between prevalence and parasite population size. Despite prolonged mass drug administration (MDA), snail control, and sanitation improvements [9,10], parasite populations retained substantial diversity, showed no detectable recent decline in effective population size, and maintained connectivity across multiple scales.

### Demographic stability despite sustained control

One explanation for the lack of demographic collapse is continued reproduction in hosts not fully captured by human prevalence metrics. *S. japonicum* infects multiple mammalian hosts, including agricultural animals such as bovines and pigs, and commensal species such as dogs, cats, and rodents [9,43–46]. Although relative contributions from these host may vary, evidence from eastern China suggests bovines likely play a major role in maintaining local parasite reservoirs, and sustaining a major fraction of human transmission [46,47]. In multi-host systems, effective population size reflects reproduction across the broader transmission network. Stable N_e_ despite declining human prevalence therefore implies continued stable reproduction within the wider host community. We do not interpret these findings as evidence that control failed. Rather, they suggest that the measured reductions in human prevalence reflect a reshaping of the overall transmission structure, allowing N_e_ to remain above maintenance levels. These findings have crucial implications for elimination strategies that rely primarily on human prevalence as an indicator of interruption.

### Localized transmission and clonal amplification within villages

Although regional effective population size remained stable, genomic patterns at finer scales revealed highly localized transmission. Within hosts, dense clusters of full- and half-siblings reflect repeated exposure to clonal cercariae produced by infected snails. Blocks of relatedness and host-level clustering (Fig 5) illustrate how asexual amplification can generate bursts of genetically identical parasites infecting multiple individuals within a village. At the village scale, broader clusters of first- and second-degree relatedness among hosts suggest a relatively high frequency of human hosts being infected by clonal miracidia from a single snail, indicated focal transmission sustained by limited snail habitats and/or abundance. Some villages were dominated by a small number of parasite lineages, producing elevated inbreeding and tightly bounded relatedness clusters, whereas others showed broader distributions consistent with more heterogeneous local parasite pools. These contrasting patterns suggest that control fragmented transmission into a limited number of local lineages rather than eliminating it entirely.

Clonal expansion in snails amplifies the effect of infections in mammals, leading to concentrated village-level transmission and persistence even when regional transmission intensity is low. Importantly, elevated inbreeding and strong within-village relatedness do not imply demographic collapse; instead, they are consistent with small but persistent transmission units embedded within a larger regional population. This mosaic structure of localized bottlenecks nested within a stable metapopulation is consistent with host–parasite metapopulation theory [19,48]. In such systems, patchy local extirpation and recolonization can occur without regional or global elimination. Such recolonization may also be driven by the long-lived nature of worms in human hosts [49],and the potential mobility of hosts over such long periods enabling the re-seeding of local infection sources. Our results suggest that transmission in Sichuan during this near-elimination period reflected exactly this combination of patchy local persistence and regional buffering.

### Limited recent dispersal but persistent regional connectivity

First- and second-degree relationships were overwhelmingly confined within villages, indicating that most recent transmission events are highly localized. Host-level networks (Fig 5A) show only rare cross-village close-kin connections, suggesting that recent dispersal between villages was uncommon, but frequent enough to be consistent with a mechanism for regional transmission. The R-P-U cluster is the clearest example of such a cross-village connection, but appears to be an exception rather than being typical. These rare cross-village links likely reflect human or human-mediated movement, or nearby reservoir-mediated connections, that bridge otherwise localized parasite populations. Modeling studies support this, demonstrating that hydrologic and social connectivity can sustain transmission cycles at low prevalence in spatially clustered systems [21,50,51].

The RAB heatmap (Fig 4) and third-degree network analyses (Fig 6B) also reveal evidence of broader regional connectivity. Moderate-relatedness signals and third-degree links spanning geographic clusters indicate that parasite lineages are not confined to single villages over longer timescales. These intermediate connections likely reflect historical or indirect transmission pathways, including low-frequency human or animal movement and hydrologically connected snail habitats. The spatial pattern of third-degree relatedness mirrors the weak north–south gradient observed in PCA and ADMIXTURE analyses, reinforcing the conclusion that villages remain part of a cohesive regional population. Even infrequent dispersal may be sufficient to prevent long-term genetic isolation and buffer parasite populations against local extinction.

Such a structure is consistent with metapopulation models in which local transmission foci are embedded within a weakly connected regional network [20,48]. In this context, elimination efforts may reduce local transmission intensity without fully disrupting regional connectivity, allowing persistence below apparent epidemiological thresholds.

### Heterogeneity in worm burden as an indicator of exposure intensity

Our analyses suggest that heterogeneity in worm burden may further shape fine-scale transmission dynamics. Using sibling clusters as proxies the minimum number of distinct reproducing worm pairs per host, we observed substantial variation in minimum worm burden across hosts and villages. Some hosts harbored parasites derived from a single adult worm pair despite multiple sampled miracidia, whereas others contained numerous genetically distinct sibling clusters, indicating multiple concurrent worm pairs and, in some cases, over 20 unique infection events, cautioning that these are a lower bound estimates due to incomplete sampling.

A minority of hosts appeared to have large numbers of adult worm pairs, consistent with well-documented individual heterogeneity in transmission [16,52] and with the classic aggregation of worm burden in schistosomiasis [13]. We do not interpret this as definitive evidence of superspreading [16], but rather it is evidence that some hosts are acquiring infections via a substantial number of unique infection events which could be due to high exposure patterns or variations in host susceptibility to infection [53]. In near-elimination settings, such heterogeneity may be particularly consequential as hosts with a large number of genetically distinct worms indicate multiple infection sources.

Extending this use of genomic data to estimate the number of unique adult worm pairs at the village level shows how genomic metrics can play a key role in guiding elimination. For example, villages S and B have comparable infection prevalence (10.4% and 11.6%, respectively), and similar sampling depth (in both villages miracidia were collected and whole genome sequenced from 3 hosts including 29 miracidia from village S, and 37 from village B). However, in village S, we inferred 6 adult worm clusters and in village B we inferred 22 This suggests that residents in village S were exposed to greater numbers of infection sources than residents in village B. This shows how simple measures of infection prevalence may obscure differences in the force of infection across villages. Integrating worm-burden heterogeneity with relatedness-based transmission mapping may therefore help prioritize focal interventions during late-stage control.

### Environmental persistence and incomplete disruption of transmission pathways

The localized clustering of closely related parasites within villages, together with rare cross-village links, implicates environmental persistence as a key component of continued transmission. *Schistosoma japonicum* depends on *Oncomelania* snails, whose distribution is shaped by hydrology, agriculture, and landscape structure [21,54]. Water networks that sustain *Oncomelania* habitats and concentrate human and livestock activity likely serve as primary conduits of parasite movement between neighboring transmission sites [18]. Even under sustained snail control, small residual habitats can maintain infected snail populations capable of generating focal outbreaks through clonal cercarial amplification.

The tightly bounded within-village relatedness patterns we observed are consistent with transmission maintained by limited infected snail foci. At the same time, moderate off-diagonal relatedness and third-degree links spanning geographic clusters suggest that environmental connectivity via waterways, seasonal flooding, agricultural water management, or human and livestock movement has not been fully disrupted. Modeling studies demonstrate that even weak spatial coupling can sustain parasite persistence below classical density-dependent breakpoints [21,50,51]. These findings do not imply widespread uncontrolled transmission.

Rather, elimination efforts appear to have fragmented transmission into localized foci without fully severing ecological and spatial pathways necessary for long-term persistence. In such contexts, re-emergence may arise from expansion of environmentally buffered transmission units within a weakly connected regional network. Targeting residual environmental linkages may therefore be as important as continued chemotherapy in late-stage elimination.

Integrating genomic surveillance with fine-scale ecological mapping could help identify persistent transmission corridors that prevalence monitoring alone may miss.

Building on the multi-scale genomic evidence presented above, particularly the absence of demographic collapse and the persistence of regional connectivity, the particular case of Sichuan provides unique additional insight into what happened next. In a prior study, Shortt et al. [38] used reduced-representation genomic data to estimate patterns of relatedness and transmission across human hosts in the years following re-emergence (2008-2016). Their results revealed stronger spatial structuring than observed in our 2007 data, with parasite genetic structure more clearly differentiated among villages and transmission largely attributable to local (e.g., within-village) reservoirs rather than widespread regional mixing. This increased structuring suggests that continued control efforts did disrupt most broader-scale transmission pathways over time. However, Shortt et al. [38] also found that parasites within villages remained highly related and that infections were frequently derived from a limited number of local sources, indicating persistent local reservoirs and transmission. Taken together with our findings from 2007, these results suggest a shift from an initially well-connected regional transmission network to a more spatially fragmented system under continued control, in which long-distance dispersal is reduced but local transmission networks remain intact. Thus, while subsequent control efforts appear to have been effective at breaking regional connectivity, they fell short of eliminating locally sustained parasite populations that continued to drive local transmission and enable persistence. From a control perspective, these results highlight a key limitation of late-stage elimination programs: disrupting regional transmission alone is insufficient if persistent local reservoirs are not identified and fully eliminated.

Achieving durable elimination will likely require targeted strategies focused on detecting and interrupting transmission within villages, confirming clearance of infection in human hosts, and addressing contributions from non-human reservoir hosts.

### Implications for elimination strategy and genomic surveillance

China’s schistosomiasis control program remains among the most successful globally, achieving dramatic reductions in morbidity and human infection prevalence over the past several decades [7,10]. Our findings do not diminish these successes. Rather, they highlight the biological complexity that can persist in near-elimination settings, where transmission operates across fragmented yet connected ecological landscapes. A central insight from this retrospective analysis is that declining human case counts do not necessarily reflect proportional reductions in parasite effective population size or transmission potential. In multi-host, environmentally mediated systems such as *S. japonicum*, persistence can be sustained through localized clonal amplification, worm-burden heterogeneity, host preference switching , and low-frequency regional connectivity. These interacting mechanisms may maintain parasite populations below apparent epidemiological thresholds, permitting re-emergence when ecological or programmatic conditions shift.

Traditional surveillance tools such as prevalence surveys, egg counts, and sentinel snail monitoring remain essential, but they primarily capture current infection status rather than underlying transmission structure. Population genomic data, by contrast, quantifies relatedness, connectivity, demographic stability, and signatures of historical patterns. In this study, genomics revealed regional cohesion, stable effective population size, dense within-village sibling clustering, and persistent inter-village connectivity – patterns not evident from prevalence data alone. As elimination programs advance globally, particularly in zoonotic or environmentally complex systems, integrating genomic surveillance into monitoring frameworks may improve detection of hidden persistence and transmission corridors [18,55]. In Sichuan, parasites collected in 2007, the year following documented re-emergence, already exhibited genomic signatures of regional stability and multi-scale connectivity. These patterns suggest that conditions permitting resurgence were present before epidemiological signals became apparent. Earlier detection of such structural persistence could support more targeted and durable elimination strategies.

## Data availability

Whole genome sequencing data generated during this work have been deposited in the NCBI Sequence Read Archive under BioProject PRJNA1365779.

## Acknowledgments

We thank the staff of the Sichuan Centers for Disease Control and the local county anti-schistosomiasis stations for their assistance with fieldwork, sample collection, and associated epidemiological data. This work was supported by the National Institutes of Health (R21 AI115288 from National Institute of Allergy and Infectious Disease and R01 AI134673-01) awarded to E.J.C. as Principal Investigator, with Y.L., B.Z., T.A.C., and D.D.P. as co-investigators. We also thank Zibiao Gou and the North Texas Genome Center for their contributions to sequencing support and computational resource access, respectively.

## Author contributions

Conceptualization: TAC, EJC, DDP, HDG Data curation: YL, BZ, EG, HG, YZF, HP, KJW Formal analysis: HG, YZF, EG, AH, WZ Funding acquisition: EJC, TAC, DDP Supervision: EJC, TAC, DDP Validation: EJC, TAC, DDP, HP, HG, YZF, EG, WZ Writing – original draft: HG, TAC, YZF, EJC Writing – review & editing: HG, YZF, EG, AH, WZ, KJW, HP, SSG, YL, BZ, DDP, EJC, TAC

## Supporting information

**S1 Table. Distribution of parasite samples and relatedness within hosts across villages.**

The estimated prevalence of infections and mean village infection intensity in 2007 for each village included in this study. Prevalence and infection intensity estimates are based on methods described in [1].

**S1 Fig. Sequencing quality metrics across 270 *S. japonicum* miracidia samples.**

Top panel shows the distribution of the percentage of reads mapped to the *S. japonicum* reference genome. Most samples exhibited high mapping rates (>95%), indicating strong enrichment for parasite DNA and minimal host contamination. Bottom panel shows the distribution of mean genome-wide sequencing coverage per sample. Coverage was generally centered between ∼20× and 35×, with a small number of higher-coverage outliers. Together, these results demonstrate consistently high-quality whole-genome sequencing suitable for downstream population genomic and relatedness analyses.

**S2 Fig. Population structure inferred using ADMIXTURE.** Cross-validation (CV) error values across K = 1–12 indicate minimal support for additional population structure beyond K = 1. Bar plots show individual ancestry proportions for K = 2 and K = 3 based on a 10-kb thinned genome-wide SNP dataset. Each vertical bar represents a single miracidium, with colors indicating inferred ancestry components. Individuals are ordered by village following a north–south geographic gradient, with village identity indicated by the color bar below. At K = 2, parasites form two broadly distributed ancestry components with limited geographic structuring. At K = 3, additional subdivision emerges, though patterns remain diffuse and do not correspond strongly to geographic groupings, consistent with weak population structure and ongoing connectivity across villages.

**S3 Fig. Concordance between RAB and RAS relatedness metrics and pedigree-based classifications.**

(A) Scatterplot of pairwise relatedness estimates comparing RAB (x-axis) and RAS (y-axis) across all pairs of *S. japonicum* miracidia. The dashed line indicates the fitted linear relationship (y = 0.592x), with R² = 0.364, demonstrating a strong positive correlation between the two metrics.

(B) Same comparison colored by Sequoia-inferred top relationship class (FS = full siblings; HA = half siblings; U = unrelated). Full- and half-sibling pairs cluster at higher RAB and RAS values, consistent with increasing genetic relatedness. (C) Same comparison colored by COLONY full-sibling assignments (probability ≥ 1), showing substantial concordance between COLONY full-sibling calls and elevated RAB/RAS values. Together, these results demonstrate agreement between likelihood-based relatedness metrics and pedigree reconstruction approaches.

**S4 Fig. Village-level network of third-degree genetic relationships across Sichuan**

Circular network (circos) plot summarizing the number and degree of pairwise genetic relationships among villages. Each outer segment represents a village, ordered geographically, with the outermost colored band indicating geographic group (North, Central, Southcentral, South). Chords connecting villages represent inferred relatedness links between miracidia sampled from different villages, colored by degree of relatedness (1st-degree, 2nd-degree, 3rd-degree). Link width is proportional to the number of inferred relationships between village pairs. First- and second-degree connections are rare and largely localized, whereas third-degree links are widespread and form a dense, regionally connected network spanning multiple geographic groups.

**S5 Fig. Distribution of parasite samples and relatedness within hosts across villages.**

Bars show the total number of *Schistosoma japonicum* miracidia sampled from each host, while black points indicate the number of observed related pairs (RAB ≥ 0.4) detected among parasites within that host. Hosts are ordered geographically from north to south by village to highlight regional patterns in parasite sampling and relatedness structure. Bar colors indicate village of origin, and the colored strip along the bottom denotes broader geographic clusters (North, Central, SouthCentral, and South). Variation in both sample counts and the number of related parasite pairs among hosts reflects heterogeneity in inferred worm burdens and transmission structure across the study region.

